## Supplementary figures and images for "Interleukin-4 induces CD11c^+^ microglia leading to amelioration of neuropathic pain in mice"

### Figure1-figure supplement 1

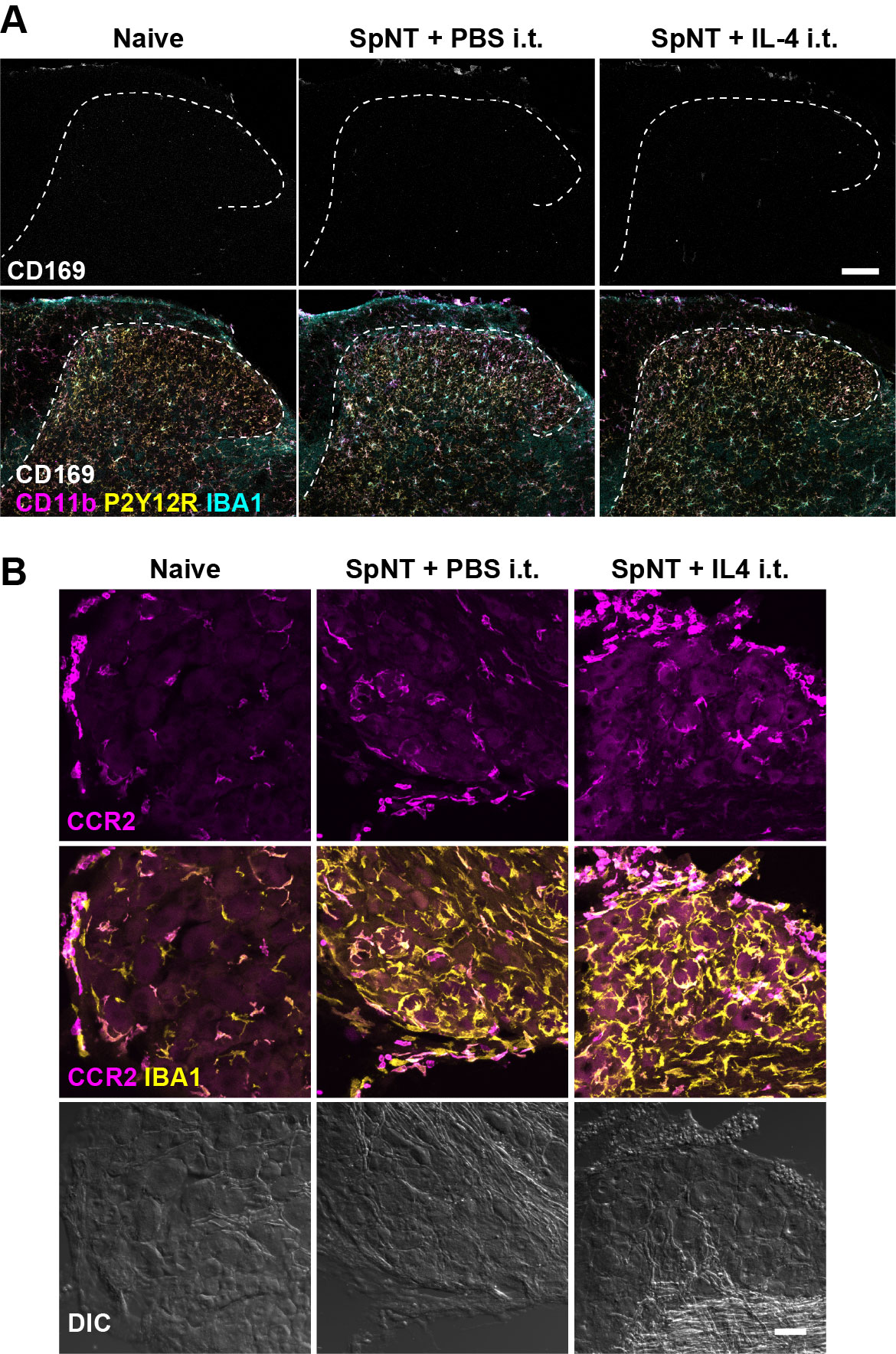
